# Valence-partitioned learning signals drive choice behavior and phenomenal subjective experience in humans

**DOI:** 10.1101/2023.03.17.533213

**Authors:** L. Paul Sands, Angela Jiang, Rachel E. Jones, Jonathan D. Trattner, Kenneth T. Kishida

**Affiliations:** Dept. of Physiology and Pharmacology, Wake Forest School of Medicine, Winston-Salem NC, 27101, US; Neuroscience Graduate Program, Wake Forest School of Medicine, Winston-Salem NC, 27101, US; Dept. of Neurosurgery, Wake Forest School of Medicine, Winston-Salem NC, 27101, US

**Author notes:** **CORRESPONDING AUTHOR** Kenneth T. Kishida, PhD; Wake Forest School of Medicine, Medical Center Blvd, Winston-Salem NC, 27101, US. Fralin Biomedical Research Institute at Virginia Tech (L. Paul Sands).

**Keywords:** consciousness, subjective experience, decision-making, reinforcement learning, reward prediction errors, punishment prediction errors, valence, affect

## Abstract

How the human brain generates conscious phenomenal experience is a fundamental problem. In particular, it is unknown how variable and dynamic changes in subjective affect are driven by interactions with objective phenomena. We hypothesize a neurocomputational mechanism that generates valence-specific learning signals associated with ‘what it is like’ to be rewarded or punished. Our hypothesized model maintains a partition between appetitive and aversive information while generating independent and parallel reward and punishment learning signals. This valence-partitioned reinforcement learning (VPRL) model and its associated learning signals are shown to predict dynamic changes in 1) human choice behavior, 2) phenomenal subjective experience, and 3) BOLD-imaging responses that implicate a network of regions that process appetitive and aversive information that converge on the ventral striatum and ventromedial prefrontal cortex during moments of introspection. Our results demonstrate the utility of valence-partitioned reinforcement learning as a neurocomputational basis for investigating mechanisms that may drive conscious experience.

**Highlights:**

- TD-Reinforcement Learning (RL) theory interprets punishments relative to rewards.
- Environmentally, appetitive and aversive events are statistically independent.
- Valence-partitioned RL (VPRL) processes reward and punishment independently.
- We show VPRL better accounts for human choice behavior and associated BOLD activity.
- VPRL signals predict dynamic changes in human subjective experience.

## INTRODUCTION

The mechanisms by which the human brain generates the subjective phenomenal experiences that allow us to answer introspective questions like, “What is it like to be [me]?” (or “a bat”; Nagel, 1974) remain a fundamental mystery that has occupied artists, philosophers, and neuroscientists for centuries (Faherty, 2016). However, this problem represents more than just an old philosophical quandary: brain states underlying subjective suffering and challenges to the ability to control one’s behavior are at the core of nearly all psychiatric and neurological conditions (Kishida et al., 2010; Montague et al., 2012; Kishida, 2012; Redish and Gordon, 2016; Huys et al., 2016; Taschereau-Dumouchel et al., 2022). The inherently subjective nature of conscious experience has led philosophers to deem an understanding of the mechanisms supporting it fundamentally ‘hard’ or even impossible (Nagel, 1974; Chalmers, 1995). On the other hand, empirical investigation has turned previously seemingly impossible problems (e.g., an understanding of electromagnetic phenomena; Forbes and Mahon, 2014) into well-defined scientific fields of inquiry. Here, “we get on with the task” of empirically investigating simple conscious experiences through the lens of behavioral and neurobiological measurements (Churchland PM, 1984, 2014; Churchland PS, 1996) that may be better understood within a neurocomputational framework (Churchland and Sejnowski, 1994; Kishida, 2012). One of the major challenges facing a science of consciousness lies in the fact that the phenomena to be investigated – e.g., variations in how one feels – are subjective and only *indirectly* accessible through self-report behavior. Nonetheless, subjective experiences are associated with reproducible behaviors and changes in neurophysiology that can be studied within behavioral, cognitive, and computational neuroscience methods.

A leading neurocomputational approach to investigating adaptive human choice behaviors and subjective experiences has been the use of temporal difference (TD) reinforcement learning (RL) theory (Sutton, 1988; Sutton and Barto, 1998) to provide a framework for probing how dopamine neurons encode ‘teaching signals’ in the form of TD reward prediction errors (RPEs; Montague et al.,1996; Schultz et al., 1997; Bayer and Glimcher, 2005; Bayer et al., 2007; Zaghloul et al., 2009; Glimcher, 2011; Hart et al., 2014; Eshel et al., 2015; Kishida et al., 2016; Watanabe-Uchida et al., 2017; Moran et al., 2018). In the dopamine TD-RPE hypothesis (Montague et al., 1996; Schultz et al., 1997), phasic bursts and pauses in dopamine neuron firing activity signal ‘better-than-expected’ or ‘worse-than-expected’ prediction errors, respectively, which provides a computationally-optimal method for learning – directly from experience – value associations between rewards and the stimuli and actions that predict them (Sutton and Barto, 1998). This mechanistic insight has since led to specific hypotheses about the neurochemical basis of computations underlying human choice behaviors and a variety of mental health disorders (Redish and Gordon, 2016), in part due to support from human fMRI studies demonstrating that BOLD activity in brain regions rich in dopaminergic terminals parametrically tracks reward prediction errors during classical and instrumental conditioning (O’Doherty et al., 2003; McClure et al., 2003; Pessiglione et al., 2006; Garrison et al., 2013). Furthermore, empirical studies have begun to associate neurocomputational processes underlying RPE encoding with the immediate subjective experience of pleasure as well as associated dynamic changes in mood that occur over longer timescales (Delgado et al., 2006; *X*iang et al., 2013; Rutledge et al., 2014; Eldar et al., 2016). This work has also provided a basis for investigating the neural and behavioral correlates of changes associated with various psychiatric conditions and mood disorders (Redish and Gordon, 2016; Redish, 2004; Montague et al., 2012; Kishida et al., 2010; Huys et al., 2016; Rutledge et al., 2017; Brown et al., 2021).

Despite its overwhelming utility, RL theory does not explicitly describe how biological organisms learn optimally from aversive experiences (i.e., punishments) concurrently or independently from experienced rewards (Sutton, 1988; Sutton and Barto, 1998; Dayan and Niv, 2008; Pessiglione and Delgado, 2015; Kishida and Sands, 2021). Aversive stimuli (e.g. those that cause tissue damage or threaten to do so) are evolutionarily conserved drivers of defensive and other adaptive behaviors (Cisek, 2021) and negative aspects of human subjective experience (Seymour et al., 2007b; Kishida and Sands, 2021). Typically, TDRL theory treats aversive experiences (e.g., punishments or costs) as ‘negative rewards’ and thus colinear with rewards along a single valence dimension: TDRL treats punishing (i.e., aversive) outcomes only *in relation to* appetitive or rewarding experiences. This is counter to biological experience where appetitive and aversive experiences may largely be derived from statistically independent sources or derived from a similar source with varying degrees of statistical dependence. For example, individual people may be collaborative or fiercely competitive; potential sources of food (e.g., plants) may be nutritious or toxic. Further, computationally, using a single valence dimension ordains a ‘zero-sum rule’ for how to represent co-occurring positive and negative outcomes (or mixed-valence experiences) – only a resultant single scalar ‘reward’ term is used in TDRL (Sutton and Barto, 1998) – and thereby also does not allow for dissociating the individual effects of co-occurring rewards and punishments on agent behavior and subjective experiences. Indeed, this traditional reliance on a unidimensional, colinear valence representation belies the independent influences of rewards and punishments (or otherwise appetitive and aversive events) on human choices and emotional responses (Konorski, 1967; Dickinson and Dearing, 1979; Cacioppo et al., 1999; Folkman and Moskowitz, 2000; Larsen et al., 2009; Larsen and McGraw, 2011; Kishida and Sands, 2021).

Fundamentally, there remains a gap in the literature regarding traditional TDRL accounts of reward and punishment learning and comparisons to alternative models that directly address how punishing stimuli may be processed in a comparably optimal manner. Human fMRI investigations of aversive valence-processing suggest that adaptive learning from punishments (e.g., pain) is consistent with a hypothetical punishment-based (i.e., reward-opponent) RL system (Palminteri and Pessiglione, 2017; Seymour et al., 2004, 2005, 2007a, 2012; Delgado, et al. 2008, 2009). Theoretical descriptions of a system ‘opponent’ to dopaminergic reward processing have been hypothesized (Daw et al., 2002) and are supported by indirect evidence (Palminteri and Pessiglione, 2017; Seymour et al., 2005, 2007a, 2007b; Delgado, et al. 2008, 2009) and recent direct simultaneous measurements of serotonin and dopamine in human striatum (Kishida et al., 2016; Moran et al., 2018). However, these prior investigations generally used a unidimensional representation of valence. To begin to explicitly compare alternatives to unidimensional TDRL-based depictions of reward and punishment learning, we hypothesized *valence-partitioned reinforcement learning* (VPRL; Kishida and Sands, 2021), which proposes that separate neural systems implement TD learning over appetitive and aversive experiences in parallel and thereby independently update representations of positive and negative expected state-action values, respectively. VPRL-encoded signals can then be operated on (e.g., integrated) or processed independently as necessary for guiding behavior, including when introspecting or reporting about one’s subjective feelings (Kishida and Sands, 2021).

Here, we test the hypothesis that VPRL is a better model than traditional TDRL for investigating human learning and decision-making behavior, (2) associated neural activity, and (3) dynamic moment-to-moment changes in subjective experience in humans. We combine data from two experiments involving human participants (N=47 total) scanned with fMRI while performing a probabilistic reward and punishment (PRP) task that uses monetary gains and losses as reinforcement (**Figure 1**A; Methods). We show that VPRL better explains participant choice behavior compared to traditional TDRL and that VPRL model parameters fit to participant choices are consistent with humans learning from rewards and punishments independently and asymmetrically (**Figure 2**). Further, we investigate the connection between VPRL learning signals and participants’ self-reported subjective feelings about received outcomes throughout the PRP task, demonstrating that the expected value of a chosen action and prediction errors over action-contingent rewards and punishments all uniquely influence – and together predict – subjective feelings about experienced outcomes (**Figure 3**). Model-based fMRI analyses reveal blood-oxygen-level-dependent (BOLD) signals that parametrically track VPRL learning signals and associated subjective feelings within a distributed network of striatal, cingulate, insular, and prefrontal regions (**Figure 4**,**5**). Our results support the notion of valence partitioning as a mechanism in the human brain for processing appetitive and aversive stimuli via independent and parallel TDRL mechanisms, which together provide more robust representations of independent appetitive and aversive value estimates in uncertain contexts. Further, our results demonstrate and we discuss the implications of VPRL as a valid neurocomputational framework for investigating the neural mechanisms underlying the dynamics of subjective phenomenal experience and associated behaviors in humans.

**Figure 1.**
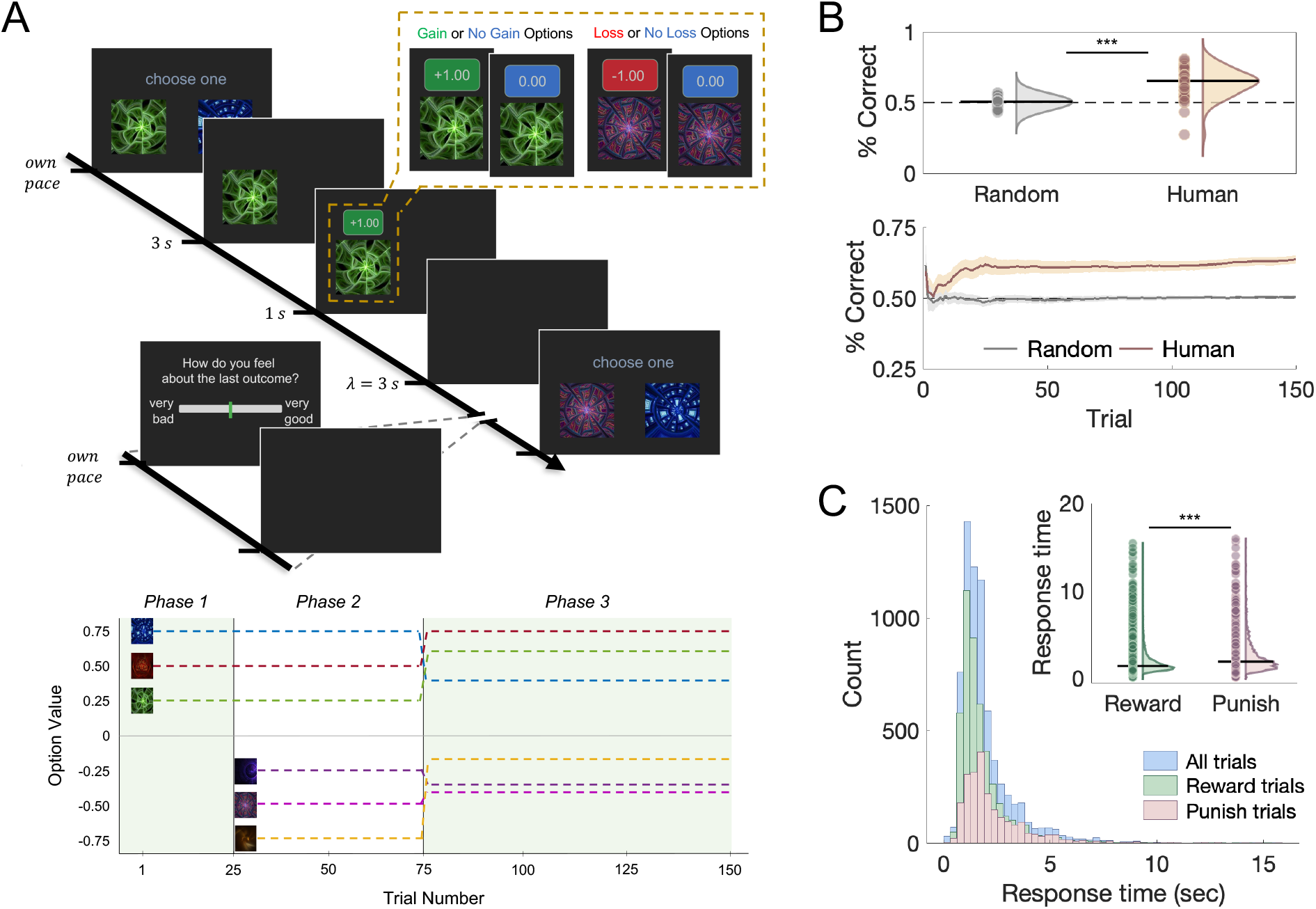
Human performance on a probabilistic reward and punishment (PRP) task. (**A**, top) Schematic of a trial from the PRP task and subjective rating prompt. On each trial, a participant chooses one of two options and is reinforced probabilistically with either a monetary gain, nothing, or a monetary loss. Randomly after a third of trials, participants submit ratings of their subjective feelings about experienced outcomes. Offset: reward-associated options result in either monetary gains or nothing, and punishment-associated options result in monetary losses or nothing. (**A**, bottom) Depiction of the ‘ground-truth’ expected value for each option (expected value = probability(outcome)*outcome) and how the options’ expected values change throughout the three phases of the PRP task (demarcated by vertical black lines). In phase 1 you choose between two of 3 possible gain/no-gain options. For phase 2, there’s an equal number of trials with two gain/no-gain options and two loss/no-loss. In phase 3, participants choose between any two of the six options at random, and the expected value for each option changes. Icon to outcome mappings are randomized for each participant. (**B**, top) The overall percent of trials where participants correctly chose the option with the highest (most positive) expected value, and (**B**, bottom) the evolution of the percent of correct choice trials throughout the PRP task. (**C**) The distributions of response times for all trials, trials on which a reward-associated option was chosen (reward trials), and trials on which a punishment-associated option was chosen (punish trials). For (**B**) and (**C**), *** = p<0.001.

**Figure 2.**
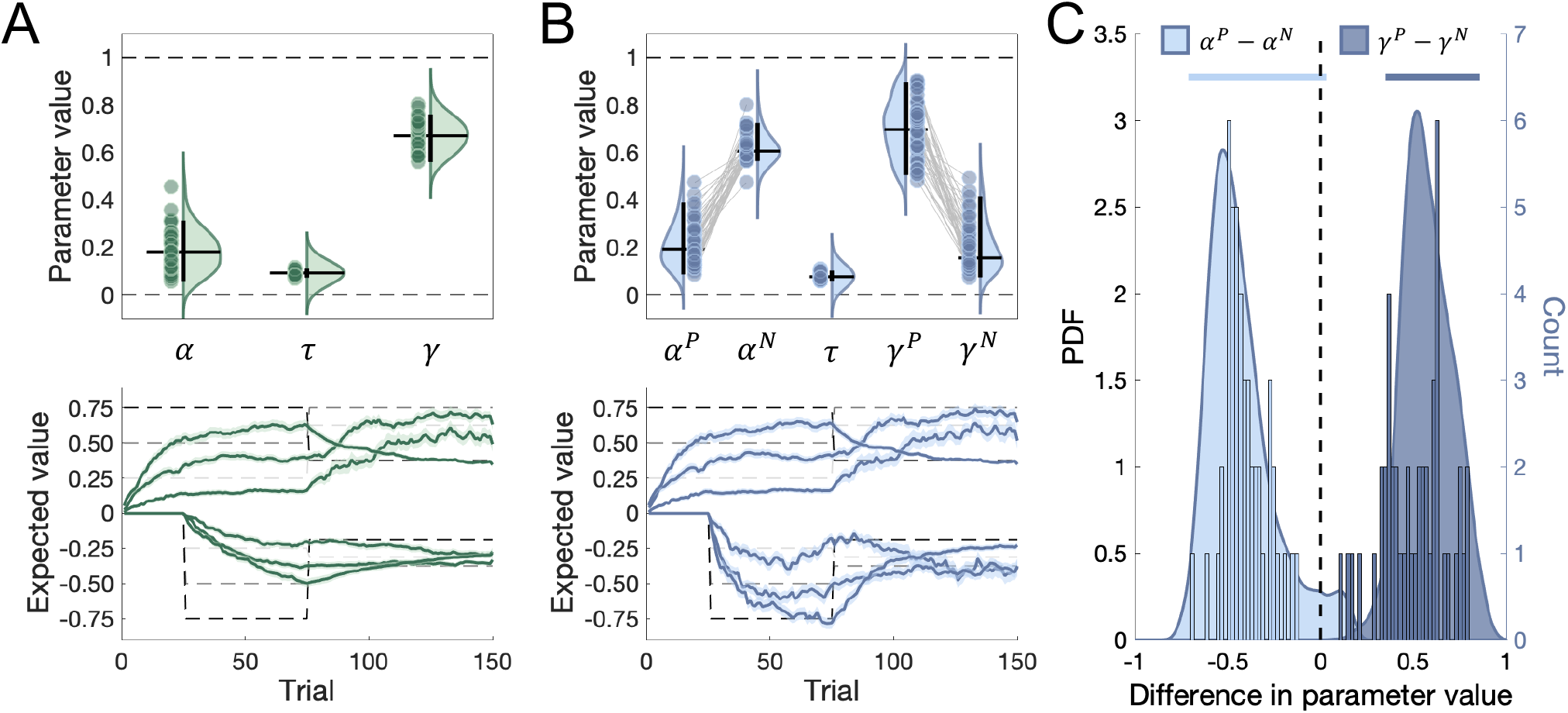
TDRL and VPRL computational modeling results. VPRL model best explains choice behavior on PRP task and leads to asymmetric learning. Distributions of (**A**) TDRL and (**B**) VPRL model parameter values across all participants. Horizontal bars mark the median of each distribution, and vertical bars indicate 95% credible interval (CI) of individual-level distribution. Dots indicate individual participant parameter values (mean of posterior parameter distributions); within-subject VPRL model parameters values are linked by grey lines. For both (**A**) and (**B**), bottom panels show time series of learned expected state-action values (Q-values) across participants for each option on the PRP task as predicated by the (**A**) TDRL and (**B**) VPRL models and shown relative to each option’s true expected value (grey dashed lines). Shaded ribbons indicate +/- one standard error of the mean (SEM), and different hues of shaded ribbons indicate different outcome probability groups. (**C**) Group- and individual-level differences in VPRL model learning rates (light blue distribution and histogram) and temporal discount parameters (dark blue distribution and histogram). Vertical dashed line indicates equivalence between parameter values; horizontal light and dark blue bars indicate 95% CI for the group-level distribution.

**Figure 3.**
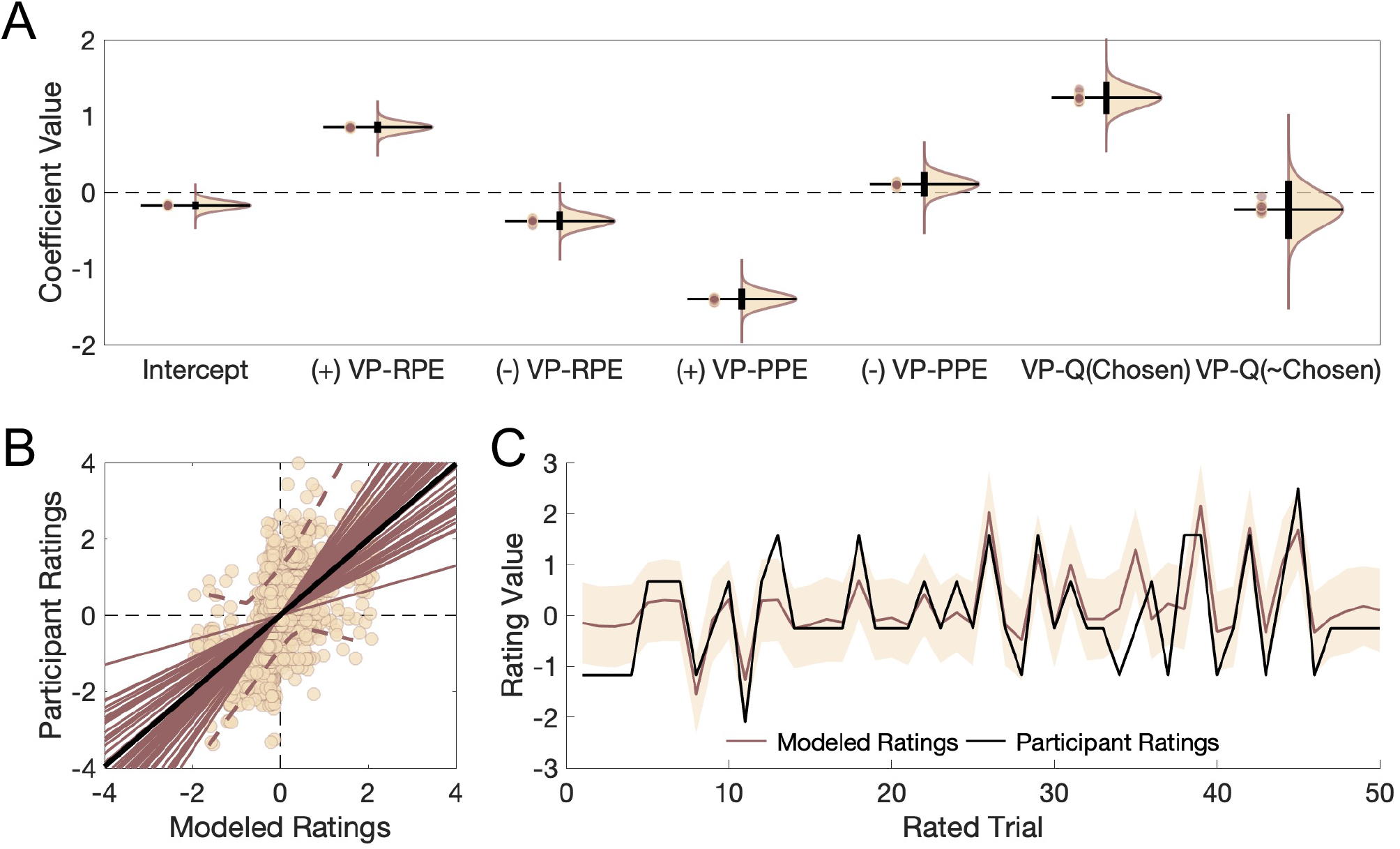
Dynamic changes in self-reported subjective feelings predicted by VPRL learning signals. Cross-validated Bayesian regression analysis reveals influence of VPRL learning signals on ratings of subjective feelings about experienced outcomes. (**A**) Distribution of regression coefficient weights on positive (+) and negative (-) VP-RPEs and VP-PPEs and learned state-action values (VP-Q-values) of the chosen and unchosen options (VP-Q(Chosen) and VP-Q(∼Chosen), respectively) on each trial. Horizontal bars indicate the median of each distribution; vertical bars indicate 95% CI of distribution; dots indicate mean individual parameter values. (**B**) Scatter plot demonstrating the linear relationship between model-derived and held-out participant ratings. Dark brown lines indicate within-participant linear correlations; black line indicates median linear relationship across all participant ratings; dashed brown lines represent 95% CI around individual-level linear correlation values. (**C**) Representative participant time series (rho=0.77, p = 7.9e-11; r-squared=0.59) of normalized subjective ratings (black line) and the cross-validated, model-predicted subjective ratings. Dark brown line represents the mean model-predicted ratings, and the shaded region represents ±1 standard deviation around mean model predictions.

**Figure 4.**
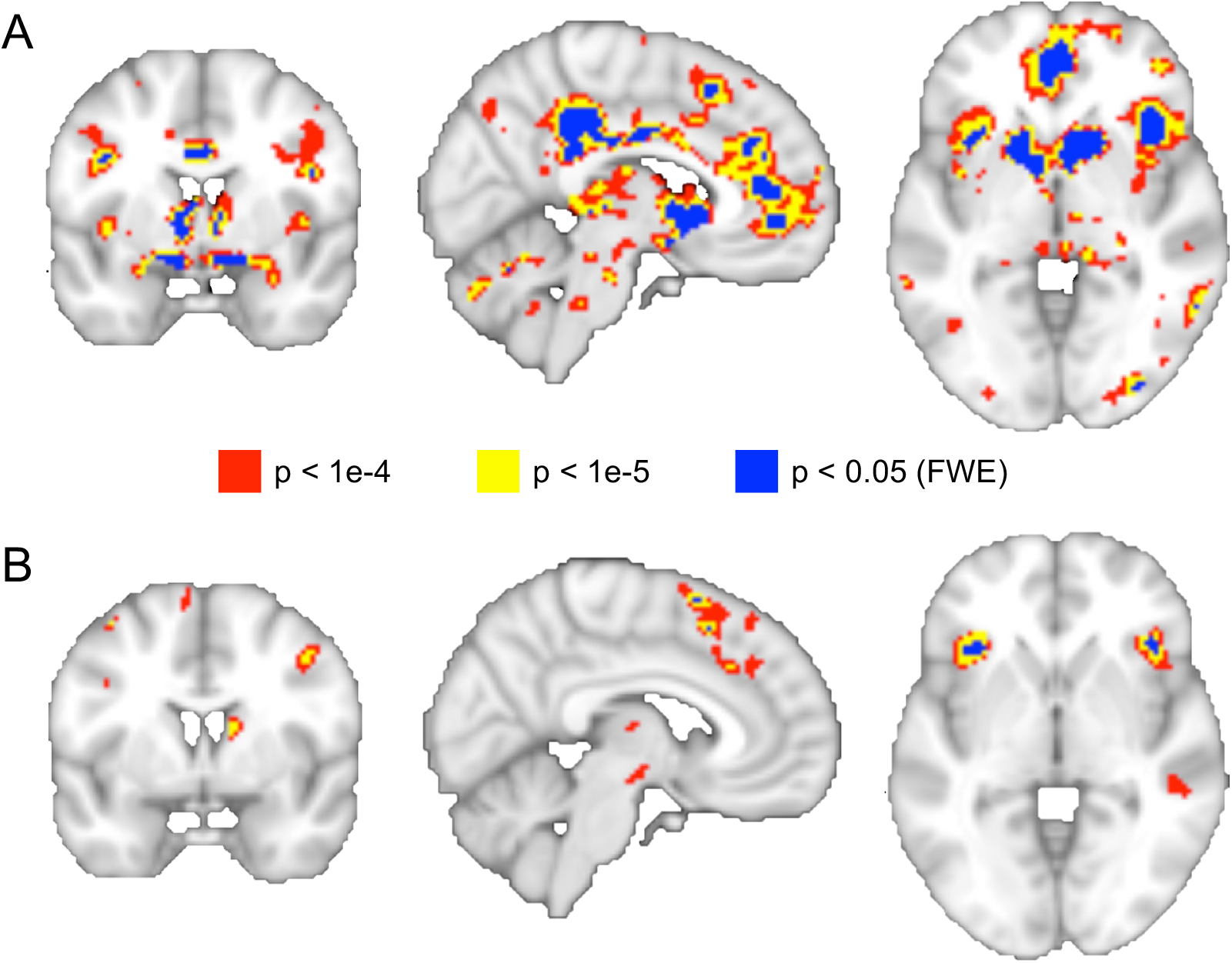
Meso-cortico-limbic regional activity represents the set of VPRL prediction error signals. Whole-brain model-based analysis of VPRL learning signals indicate that regions of human striatum, insula cortex, and anterior cingulate cortex parametrically encode (**A**) VP-RPEs or (**B**) VP-PPEs. Colored voxels and associated p-values indicate statistical thresholding used for primary analyses. All panels are sliced at MNI coordinates *X*=6, y=2, z=-2. FWE = family-wise error.

**Figure 5.**
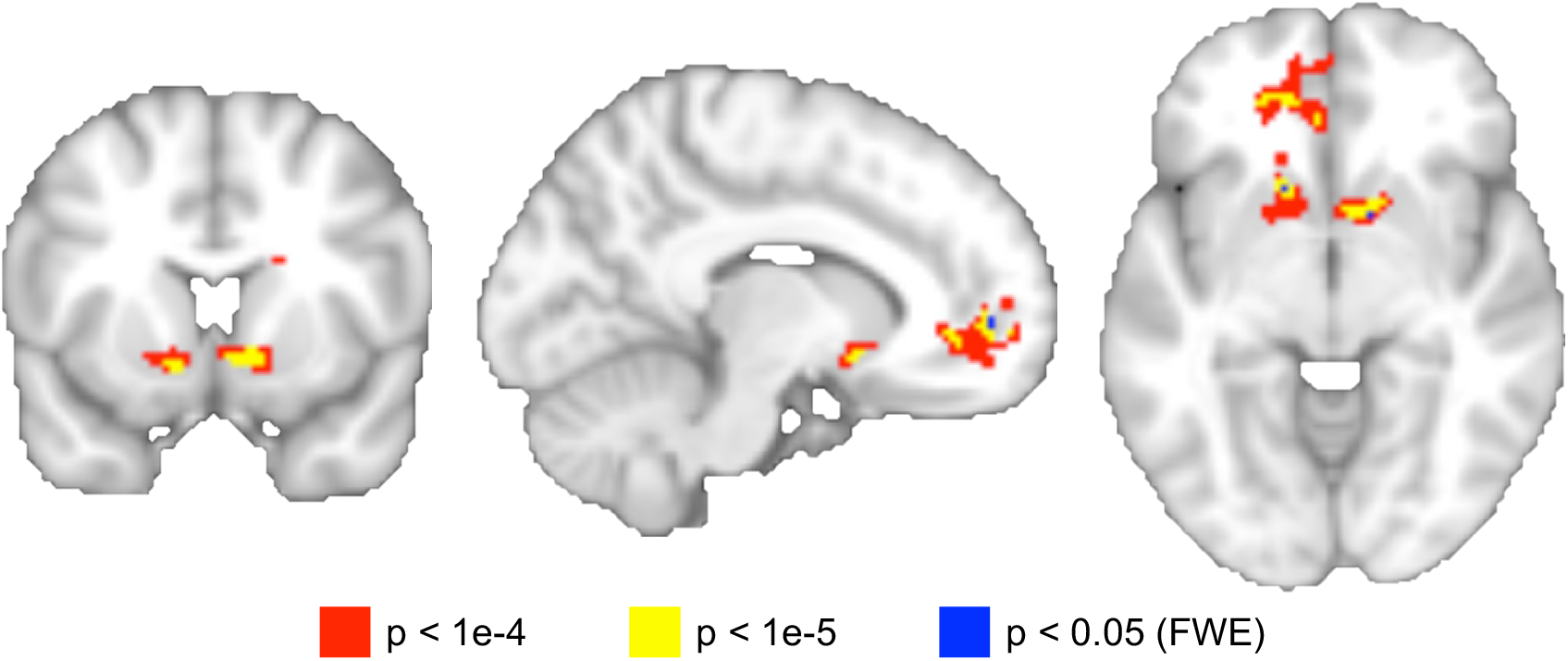
Medial prefrontal and ventral striatal activity correlates of trial-to-trial subjective human feelings. Whole-brain model-based analysis of self-reported subjective feelings as predicted by VPRL expected values and prediction errors indicate that medial prefrontal cortex and ventral striatum parametrically track the imputed subjective feeling on each trial. Colored voxels and associated p-value indicates statistical thresholding used for primary analyses; *X*=-10, y=-10, z=6. FWE = family-wise error.

## RESULTS

### VPRL best explains choice behaviors and reveals asymmetrical processing of rewards and punishments

Forty-seven participants completed the PRP task which required them to intermittently (randomly on one-third of all trials) rate how they felt about recent outcomes (**Figure 1**A). Participants learned to choose the option with the highest expected value on each trial more often than expected by chance (**Figure 1**B; two-sample t-test, t(92)=8.8, p<0.001), chose the option with the highest expected value increasingly over time (two-way ANOVA (group, time), F(1,149)=2416.8), and were quicker to select rewarding options than punishing options (**Figure 1**C; two-sample t-test, t(7023)=-14.5, p<0.001).

To test whether participants might learn differently from rewards and punishments, we fit a standard TDRL model and a VPRL model (Kishida and Sands, 2021) to participant choice behavior using hierarchical Bayesian inference and compared estimates of the model evidence (i.e., marginal likelihood) and the posterior predictive accuracy (density) for both models. For both cohorts individually, and for an ‘internal meta-analysis’ combined cohort, VPRL demonstrated both greater model evidence and greater posterior predictive accuracy relative to TDRL (**Table 1**), indicating that VPRL is a better explanation of behavior on the PRP task and better predicts unobserved PRP task choice behavior data.

**Table 1.**
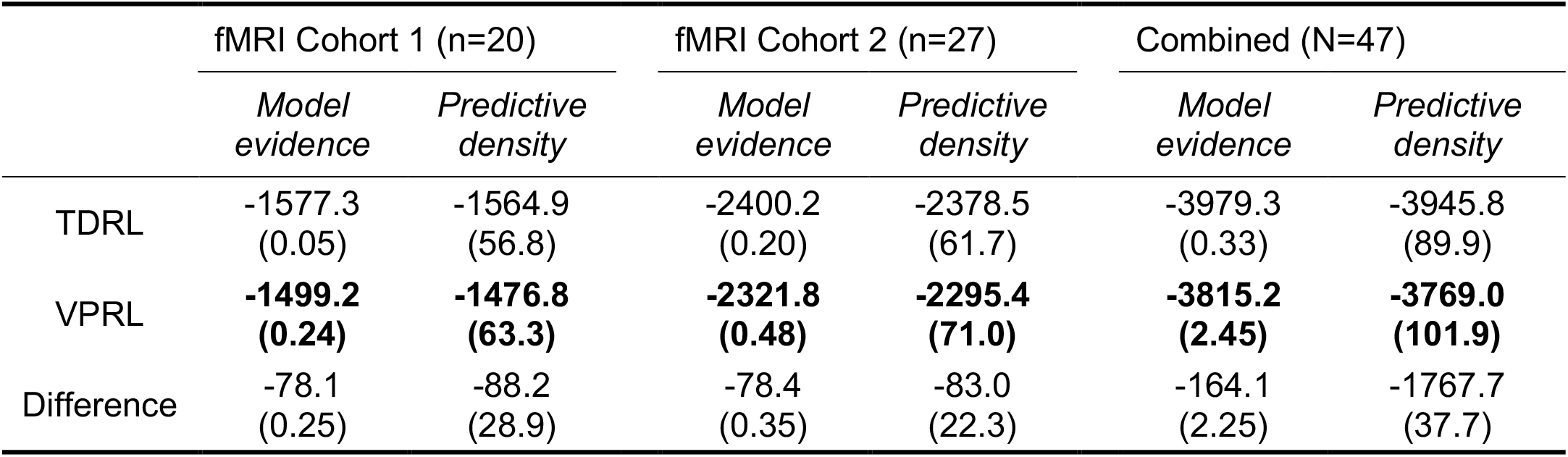
TDRL and VPRL model comparison results for two neuroimaging cohorts. Computed estimates of the Bayesian model evidence (i.e., marginal likelihood) and model predictive density (i.e., cross-validated error) for TDRL and VPRL models. VPRL demonstrated the maximum (least negative) model evidence and expected predictive density (bold values) compared to TDRL. Reported values are the median estimate value (log scale), with values in parentheses indicating either the interquartile range (of model evidence estimations) or the Monte Carlo sampling error (for the predictive density). Given the similarity of TDRL and VPRL model comparison results for both fMRI cohorts separately and the improved model evidence and predictive density for VPRL when combining both fMRI cohorts, we elected to combine the data from both fMRI cohorts to improve the power of both the main behavioral and model-based fMRI analyses.

Given that participant choice behavior on the PRP task is better explained by VPRL compared to TDRL, we next investigated differences in the (posterior) parameter distributions and the time series of learned state-action values (Q-values) derived from each model (**Figure 2**). The group-level TDRL model parameters are (**Figure 2**A): learning rate (*α*): median = 0.16 (95% credible (highest density) interval (CI) = [0.13 0.21]); temporal discount factor (γ): median = 0.65 [0.46 0.98]; and choice temperature (*τ*): median = 0.09 [0.05 0.20]. The group-level VPRL model parameters are (**Figure 2**B): <u>P</u>ositive valence (i.e., reward) system learning rate (*α*^*p*^): median = 0.20 [0.16 0.24]; <u>N</u>egative valence (i.e., punishment) system learning rate (*α*^*N*^): median = 0.66 [0.17 0.91]; Positive system temporal discount factor (*γ*^*p*^): median = 0.71 [0.54 0.99]; Negative system temporal discount factor (*γ*^*N*^): median = 0.15 [0.03 0.29]; and choice temperature (*τ*): median = 0.07 [0.05 0.14]. To investigate the nature of the differential learning of rewards relative to punishments in the VPRL framework, we assessed the difference between the Positive and Negative systems’ learning rates and temporal discount parameters (**Figure 2**C). We found that the learning rate for punishments is generally greater than the learning rate for rewards (*α*^*p*^− *α*^*N*^ median difference = -0.47 [-0.71 0.04]), and that temporal discounting for punishments was greater than temporal discounting for rewards (*γ*^*p*^− *γ*^*N*^ median difference = 0.56 [0.35 0.86]). Lastly, the time series of learned expected values for TDRL (**Figure 2**A, bottom) and VPRL (**Figure 2**B, bottom) models demonstrate that participants learned option values that recapitulate the correct ranking (i.e., from most negative to most positive value) and are appropriately adaptive to the changes in outcome magnitudes beginning in Phase 3. Of note, VPRL produced more accurate estimates of the true state-action values of aversive options (i.e., associated with monetary losses) over time compared to TDRL (**Figure S1**; two-way ANOVA (time, model): F(time,149) = 8.7, p = 3.1e-83; F(model,1) = 66.2, p = 2.4e-15), whereas rewarding options are learned with equivalent accuracy (F(time,149) = 2.38, p = 1.9e-13; F(model,1) = 0.02, p = 0.66); differences between learned option values for TDRL and VPRL models were specific to the most negatively valued options.

### VPRL prediction errors over rewards and punishments differentially affect subjective feelings

Given the evidence that VPRL best explains participant behavior on the PRP task, we sought to characterize how VPRL model-derived reward prediction errors (VP-RPE), punishment prediction errors (VP-PPE), and expected action values influence moment-to-moment changes in participants’ self-reported subjective feelings about experienced outcomes. We fit a cross-validated (leave-one-participant-out) Bayesian linear regression model to predict subjective feeling reports (**Figure 1**) using as predictor variables the learned state-action values (VP-Q-value) of each option presented on a rated trial and positive and negative VP-RPEs (from the VPRL Positive system) and VP-PPEs (from the VPRL Negative system) in response to the outcome of each rated trial (**Figure 3**A). Positive and negative VP-RPEs contribute positively and negatively to participants’ self-reported subjective feelings, respectively (positive VP-RPE: median = 0.92 [0.70 1.1]; negative VP-RPE: median = -0.33 [-0.64 -0.01]). Conversely, positive and negative VP-PPEs contribute negatively and positively to subjective feelings, respectively (positive VP-PPE: median = -1.4 [-1.8 -1.1]; negative punishment VP-PPE: median = 0.13 [0.02 0.21]). The expected value of the chosen option on rated trials contributes positively to subjective ratings (Expected Value (VP-EV) of Chosen: median = 1.4 [1.1 1.6]), whereas the expected value of the unchosen option on rated trials shows no effect (VP-EV of Unchosen: median = -0.05 [-0.18 0.09]).

To assess the posterior predictive performance of the cross-validated Bayesian regression model on held-out participant ratings, we computed r-squared values and Pearson correlation coefficients (rho value) between the held-out participant’s ratings and the model-predicted ratings (**Figure 3**B). This analysis revealed a median within-participant correlation measure of 0.65 (SD = 0.16; median p-value = 4.3e-7) and r-squared value of 0.42 (SD = 0.18), indicating that the cross-validated regression model generalizes moderately well to out-of-sample participant data (**Figure 3**C).

Further analyses comparing the coefficient values of VP-RPEs and VP-PPEs revealed that the magnitude (absolute value) of positive VP-RPE coefficients is generally larger than the magnitude of negative VP-RPE coefficients (median difference = 0.15 [-0.08 0.37]). Similarly, we found that positive VP-PPEs have a consistently larger contribution to subjective ratings than negative VP-PPEs (median difference = 0.89 [0.63 1.1]). Lastly, negative VP-PPEs consistently had a more positive contribution to subjective ratings than negative VP-RPEs (median = 0.27 [-0.003 0.55]). As a whole, comparing the effects of positive and negative VP-RPEs and VP-PPEs on subjective ratings revealed a consistent ordering of the contributions of VPRL prediction errors to subjective feelings, with positive VP-RPE > negative VP-PPE (median difference = 0.98 [0.74 1.2]), negative VP-PPE > negative VP-RPE, and negative VP-RPE > positive VP-PPE (median difference = 0.62 [0.40 0.84).

### Striatal-insular-prefrontal network activity tracks ‘reward’ and ‘punishment’ prediction errors

Based on related prior work, we hypothesized that VPRL reward prediction errors (VP-RPEs), punishment prediction errors (VP-PPEs), and the respective positive and negative system expected values would be tracked by regions of dorsal and ventral striatum, cingulate cortex, and insula (O’Doherty et al., 2003; McClure et al., 2003; Pessiglione et al., 2006; Palminteri and Pessiglione, 2017; Seymour et al., 2004, 2005, 2007a, 2012; Delgado et al., 2008, 2009; Garrison et al., 2013). Using a model-based approach, we tested for regional activity that correlated with VP-RPEs and VP-PPEs by computing contrasts for positive effects of VP-RPEs or VP-PPEs (**Figure 4**). Regions that show hemodynamic signatures of VP-RPEs include the anterior cingulate cortex (ACC), anterior insula, and ventral striatum (**Figure 4**A); regions that correlate with VP-PPEs included the ACC, anterior insula, and dorsal striatum (**Figure 4**B). Additionally, we found that regional activity in the ventromedial prefrontal cortex (vmPFC) tracked VPRL-derived learned action values (VP-Q-values) of both the chosen and unchosen option on each trial (**Figure S2**).

### Ventral striatum and ventromedial prefrontal cortex track participants’ subjective experience

Prior reports demonstrate that subjective feelings associated with prediction error and expected value signals are tracked by medial prefrontal cortex (Xiang et al., 2013) and ventral striatum (Rutledge et al., 2014). We hypothesized that the neural responses to signals derived from the VPRL model would drive brain responses associated with subjective feelings on every trial. Thus, we used the fitted model coefficients from the cross-validated subjective rating regression analysis to impute the subjective feeling on each trial conditioned on the trial’s prediction errors and expected value signals. We found that regional activity in ventral striatum and ventromedial prefrontal cortex (vmPFC) parametrically tracked the imputed subjective rating on each trial throughout the PRP task (**Figure 5**).

## DISCUSSION

Here, we investigated how the human brain learns from independent appetitive and aversive experiences to adapt choice behaviors and how this dynamic process impacts subjective experience. We hypothesized VPRL (Kishida and Sands, 2021) as a framework for studying how neural systems may process rewards statistically independently from punishments. We demonstrate that VPRL consistently explains human choice behavior better on a probabilistic reward and punishment task compared to traditional TDRL. Furthermore, we show that VPRL-derived expected action values and prediction errors predict participants’ self-reported ratings of subjective feelings to rewards and punishments trial-to-trial. Moreover, we demonstrated that these VPRL-derived learning signals are parametrically tracked by BOLD activity in dorsal and ventral striatum, cingulate cortex, anterior insula, and prefrontal cortex brain regions, and that BOLD signals in the ventral striatum and ventromedial prefrontal cortex track the expected rating of participants’ subjective experience on each trial. Together, our results provide insight into (1) the type of learning mechanisms in humans responsible for ascribing valence information to stimuli and actions based on experience, and (2) how distributed neural activity implementing such mechanisms may support the composition of subjective phenomenal experience.

Central to the VPRL hypothesis is the idea that there are parallel neural systems that process positive and negative experiences separately using TD learning before respective learning signals or valent value representations are available for further processing (Montague et al., 2016; Kishida and Sands, 2021). In this way, VPRL provides a novel computational framework for investigating a variety of neurophysiological, behavioral, and psychological phenomena. From an evolutionary perspective, early vertebrates likely developed neural circuits for detecting and escaping threatening (i.e., aversive) stimuli alongside, but separate from, (putatively) dopaminergic neural circuits for initial forms of associative reward learning (Cisek, 2021). This evolutionary theory is consistent with the idea that mammalian choice behavior may be driven by the activities of – and interaction between – separate positive and negative valence-processing systems, an idea with a venerable history in psychological theories of emotions (Cacioppo et al., 1999; Folkman and Moskowitz, 2000; Larsen et al., 2009; Larsen and McGraw, 2011) and motivated behaviors (Konorski, 1967; Dickinson and Dearing, 1979; Seymour et al., 2007b; Boureau and Dayan, 2013). Here, the VPRL framework can be viewed as an explicit generative account for the wide repertoire of ‘approach-avoid’ motivated behaviors (Dickinson and Dearing, 1979) and valence-specific affective responses (Cacioppo et al., 1999; Folkman and Moskowitz, 2000), while also providing a theoretical framework for considering the mechanisms of interaction between opponent systems that may lead to ‘freezing’, non-action responses (Boureau and Dayan, 2013), or the subjective phenomenal experience of ‘mixed’ or ‘conflicting’ emotions (Larsen and McGraw, 2011).

In line with these evolutionary and psychological theories, we hypothesize a VPRL model that accounts for the premise that costs and benefits are always intertwined for biological creatures constrained by metabolic, survival, and reproductive goals and demands (Montague and King-Casas, 2007; Botvinik et al., 2015). Importantly, VPRL specifies a different perspective on how valence, within the context of RL, is processed: aversive stimuli that are immediately (or predicted to be) costly are learned directly and independently from potential rewarding stimuli. This is distinct from the more common representations that requires aversive stimuli to be compared to ‘expectations’ and requires prediction error encoding according to the valence of the TD-RPE (i.e., differential learning from positive or negative RPEs).

Our present results using a simple probabilistic reward and punishment learning task indicate that independently processing rewards and punishments via VPRL reveals an increased sensitivity to immediate punishments compared to rewards and increased temporal discounting of future punishments compared to future rewards (**Figures 2**,**3**), which is consistent with prior behavioral observations (Kahneman and Tversky, 1979; Tom et al., 2007). This differential learning from gains versus losses within the VPRL framework reveals that participants learn expected values of reward at a similar rate to that expected via traditional unidimensional TDRL, though they learn expected values of losses at a much faster rate and with improved accuracy and precision (**Figure S1**). This observation suggests that VPRL signals may also independently and asymmetrically influence subjective feelings. Our results suggest that omitted or ‘smaller-than-expected’ rewards (i.e., negative VP-RPEs) do not contribute to what it ‘feels like’ in the same manner as ‘larger-than-expected’ punishments (positive VP-PPEs), nor do ‘smaller-than-expected’ punishments (negative VP-PPE) contribute to what it ‘feels like’ in the same manner as ‘larger-than-expected’ rewards (positive VP-RPEs). Such distinctions cannot be parsed within a unidimensional representation of valence as in the traditional TDRL framework. The presence of positive VP-RPEs had a significantly greater positive influence on subjective ratings than negative VP-PPEs; the opposite was also true: negative VP-RPEs had a consistently smaller negative influence on subjective ratings than positive VP-PPEs. Such relationships might reflect a relative scaling principle for positive and negative prediction errors over rewards and punishments in generating momentary affective subjective feelings, an effect that may be dependent on ventral striatal and ventromedial prefrontal neural activity (**Figure 5**). Regardless of the mechanisms to be discovered, our results demonstrate that VPRL is a valid neurocomputational framework for empirically investigating how complex interactions of reward and punishment may lead to self-reports about subjective phenomenal experience in humans.

Numerous investigations into the neural basis of prediction error signaling in humans, using a variety of computational models and experimental designs, implicate a distributed network of brain regions in tracking prediction errors (Garrison et al., 2013). Our model-based event-related fMRI results indicate that VPRL prediction errors over rewards and punishments are represented by partially overlapping regional activation patterns and along a ventral-dorsal axis within the striatum (**Figures 4**,**5**), which is consistent with prior work (Seymour et al., 2007a). We hypothesize that striatal, insular, cingulate, and prefrontal cortex functional interactions – driven by an underlying neural architecture that broadcasts VPRL learning signals throughout the brain – can be viewed collectively as part of a dynamic affective core that regulates behavioral control and is a core component underlying subjective phenomenal experience (Kishida and Sands, 2021). Indeed, ascending neuromodulatory systems that project throughout both subcortical and cortical brain regions are integral to coordinating systems-level functional interactions towards accomplishing or switching between cognitive or behavioral tasks (Shine et al., 2019, 2021). Along these lines, future work may address how dynamic patterns of activity within the distributed subcortical-cortical network identified in our VPRL model-based analysis forms representations of state-action-outcome associations and how they co-evolve with representations of affective subjective experiences. In this regard, our results outline a potential role for ventral striatum and ventromedial prefrontal cortex interactions in mediating experience-dependent changes in brain activity associated with dynamic changes in subjective phenomenal experience (Xiang et al., 2013; Rutledge et al, 2014; Eldar et al., 2016; Tom et al., 2007; Chang et al., 2021), consistent with the dynamic affective core hypothesis (Kishida and Sands, 2021).

Non-invasive brain activity measurements like fMRI are unable to provide information on the neurochemical basis of VPRL-reward prediction errors or VPRL-punishment prediction errors, though recent advances provide an opportunity for testing competing hypotheses (Kishida et al., 2016; Moran et al., 2018; Bang et al., 2020). Neurobiologically, an independent aversive system involved in valence processing may be implemented by a variety of possible neural substrates, such as a distinct population of dopamine neurons tuned for aversive stimuli (Matsumoto and Hikosaka, 2009; Lammel et al., 2014; Kishida et al., 2016; Kishida and Sands, 2021) or the serotonin neurotransmitter system (Daw et al., 2002; Boureau and Dayan, 2013; Montague et al., 2016; Moran et al., 2018; Kishida and Sands, 2021). For instance, direct electrochemical recordings of dopamine and serotonin microfluctuations in human striatum during a sequential investment task (Kishida et al., 2016; Moran et al., 2018) are consistent with the notion that these neurotransmitter systems can act as positive and negative valence-processing systems, respectively (Montague et al., 2016; Kishida and Sands, 2021). Further, dissociable effects of rewards and punishments on reversal learning have been linked to dopamine and serotonin transporter polymorphisms, respectively (den Ouden et al., 2013). Distributional RL (Dabney et al., 2020) has recently been demonstrated as a mechanism for representing a wide dynamic range of reward magnitude; still, it remains unclear how dopamine neurons come to achieve varying value prediction ‘set points’ as well as whether and how they might encode a distribution over aversive experiences. Alternatively, VPRL-like hypotheses for future investigation might address the potential distributional coding of rewards and punishments in candidate neuromodulatory systems (e.g., dopamine and serotonin; Montague et al., 2016; Kishida and Sands, 2021; Moran et al., 2018; Bang et al., 2020).

Human behavior and subjective self-reports about associated phenomenal experiences, good and bad, are multidimensional. Prior work investigating computational models and associated BOLD imaging measurements of brain activity associated with subjective experience, mood, and subjective well-being utilized traditional unidimensional reinforcement learning models as a framework (Delgado et al., 2006; Rutledge et al., 2014; Eldar et al., 2016) and inspired the present work. Here, however, we demonstrate that a unidimensional reward prediction error is not enough (in contrast to arguments presented in Silver et al., 2020; Vamplew et al., 2021) to fully account for the dynamics of human choice behavior and associated subjective experiences and can even be detrimental when reward-associated actions also incur substantial physical costs or other negative externalities that cannot be disentangled with traditional TDRL (Elfwing and Seymour, 2017). Instead, our results using VPRL suggest that (at least) two valence dimensions are necessary, but this is almost certainly far from a complete depiction of the generative signals involved in experiences and behaviors associated with ‘what it is like’ to be (Nagel, 1974). Our results are consistent with a need to account for appetitive and aversive input in parallel, though independently, such that the integration of these signals can be performed downstream of the systems that generate the error signals. As but one possible approach, VPRL maintains the computational advantages of TDRL, but also better accounts for information that biological agents must track (e.g., costly punishments or losses) that are often independent from co-occurring appetitive stimuli. We have taken an initial step to test VPRL as a hypothetical framework for investigating basic questions about how humans adapt their choice behavior and how associated signals may account for subjective phenomenal experiences. Our findings imply that new insights may be gained should VPRL (or other valence-partitioning models) be applied to computational psychiatric problems (Montague et al., 2012; Huys et al., 2016; Redish and Gordon, 2016; Brown et al., 2021) where subjective suffering and fundamental changes in adaptive behavior characterize severe challenges to mental health.

## ACKNOWLEDGEMENTS

The authors would like to thank Read Montague and Terry Lohrenz for comments on an earlier draft of the manuscript. This work was supported by NIH-NIMH R01MH121099 (KTK), NIH-NIDA R01DA048096 (KTK), NIH-NIMH R01MH124115 (KTK), NIH-NIDA P50DA006634 (KTK), NIH 5KL2TR001420 (KTK), NIH-NIDA F31DA053174 (LPS), and NIH-NIDA T32DA041349 (LPS). We would like to acknowledge the Translational Imaging Program (TIP) of the Wake Forest CTSI, which is supported by the National Center for Advancing Translational Sciences (NCATS), National Institutes of Health, through Grant Award Number UL1TR001420.

## AUTHOR CONTRIBUTIONS

L.P.S. – Collected data; designed and performed data analysis; interpreted results, wrote, edited manuscript drafts, and approved final manuscript.

A.J. – Coded behavioral tasks; collected data; analyzed data; edited and approved final manuscript

R.E.J. – Collected data; analyzed data; edited and approved final manuscript

J.D.T. – Analyzed data; edited and approved final manuscript

K.T.K. – Conceived the study; designed experiments; supervised and guided data collection and analysis; interpreted results, wrote, edited manuscript drafts, and approved final manuscript.

## DATA AND CODE AVAILABILITY

Anonymized individual-level participant behavioral task data and MRI data used in this study may be made available upon submission of a formal project outline from any qualified investigator to the corresponding author and subsequent approval by the corresponding author in line with data protection regulations of Wake Forest University School of Medicine Institutional Review Board (IRB). Custom-written analysis scripts for generating the behavioral and imaging results of this manuscript are maintained in a private github repository (*<u>insert link upon acceptance</u>*) that may be shared upon request from any qualified investigator to the corresponding author.

## METHODS

### Participants

A total of 47 participants (across two neuroimaging experiments (n=20; and n=27) were recruited from the local Winston-Salem community to complete the probabilistic reward and punishment (PRP) task. In the first fMRI cohort, participants (n=20; 16 female) were recruited from the community in Winston-Salem, NC. For the second fMRI cohort, participants (n=27; 19 female) were recruited as ‘control participants’ for an ongoing study. Recruitment of these participants in the second fMRI cohort was similar to the first fMRI cohort. However, consent to participate included repeated visits to be completed after an initial visit where the tasks completed include the PRP task as well as more extensive behavioral characterization after the PRP task was completed; the first visit in this ongoing study is similar to the visit completed by participants in the first fMRI cohort, except that *after* the completion of the PRP task with scanning participants underwent a more involved psychiatric evaluation process to properly control for the observational experimental group. Informed written consent was obtained from each participant, and the experiment was approved by the Institutional Review Board (IRB#’s: IRB00042265, IRB00054337, and IRB00056131) of Wake Forest University Health Sciences (WFUHS). All experiments were conducted at WFUHS.

Three participants’ subjective rating data were not used in the regression modeling analysis due to limited variability in the responses on the subjective rating assessment (i.e., choosing the same rating across 90% of rated trials). This results in n=44 participants for the combined fMRI cohort. From our leave-one-participant-out cross-validation approach, we computed Pearson’s correlation coefficient (rho) and r-squared values for the cross-validated model-predicted ratings (defined as the mean of samples of the posterior predictive density for the held-out participant’s ratings) and actual held-out participant ratings. This procedure was iterated across participants, such that each individual acted as the held-out participant once. We used the fitted subjective rating model coefficients for each participant (i.e., the mean model coefficients for the cross-validated model iteration when the participant was held out) to impute a subjective rating for all trials for that participant, which we incorporate into the participant’s first-level GLM in our model-based fMRI analysis.

### Probabilistic Reward and Punishment (PRP) task experimental procedure

The PRP task (**Figure 1c**) is a 150-trial, two-choice monetary reward and punishment learning task, where chosen options are reinforced probabilistically with either monetary gains (or no gain) or monetary losses (or no loss). Six options (represented by fractal images) comprise the set of possible actions, with each option assigned to one of three outcome probabilities (25%, 50%, and 75%) and one of two outcome valences (monetary gain or loss); the assignment of options to outcome probabilities and valences is randomized across participants. The task proceeds through three phases. At the beginning of the experiment (Phase 1, trials 1-25), each trial starts with the presentation of two of the three possible ‘gain/no gain options, and participants are reinforced with either a monetary gain or nothing ($1 or $0) according to the chosen option’s fixed probability. In Phase 2 (trials 26-75), the game introduces trials which present two of the three ‘loss/no loss’ options that result in either a monetary loss or nothing (-$1 or $0) with fixed probabilities. There are 25 ‘gain/no gain’ and 25 ‘loss/no loss’ trials randomly ordered in Phase 2. In Phase 3 (trials 76-150), two options are presented randomly such that any trial may consist of two ‘gain/no gain options, two ‘loss/no loss’ options, or one ‘gain /no gain and one ‘loss/no loss’ option. Moreover, in Phase 3 the outcome magnitudes of all options change: the 25%, 50%, or 75% ‘gain’ options now payout $2.50, $1.50, and $0.50 respectively, and the 25%, 50%, or 75% ‘loss’ options now lose - $1.25, -$0.75, and -$0.25, respectively (dashed lines in **Fig. 1**A, bottom).

A participant is presented with two options at the beginning of each trial, and they select an option at their own pace. The unchosen option disappears at the same time the chosen option is highlighted, and this screen lasts for three seconds. The outcome is then displayed for one second followed by a blank screen that lasts for a random time interval (defined by a Poison distribution with λ = 3 seconds) before the next trial begins. After each trial, with probability 0.33, the blank screen following outcome presentation is followed by a subjective feeling rating screen with the text “How do you feel about the last outcome?”. Participants are asked to rate with a visual-digital scale (**Figure 1**) their feelings about the last outcome, after which the blank screen reappears for another random interval before a new trial begins.

### Temporal Difference Reinforcement Learning (Q-learning) model

In the standard ‘unidimensional’ TDRL model (Sutton 1988; Watkins and Dayan, 1992; Sutton and Barto, 1998), the expected value of a state-action pair *Q*(*s*_*i*_, *a*), where *i* indexes discrete time points in a trial, is updated following selection of action *a* in state *s*_*i*_ according to the update rule:

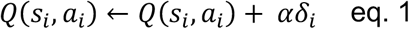

where 0 <*α* < 1 is a learning rate parameter that determines the weight prediction errors have on updating expected values, and *δ*_*i*_ is the TD reward prediction error term:

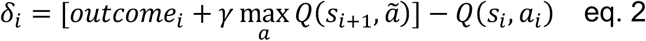

where *outcome*_*i*_ is the outcome (positive or negative) experienced in state *s*_*i*_ after taking action *a*_*i*_, 0 <*γ* < 1 is a temporal discount parameter that discounts outcomes expected in the future relative to immediate outcomes (i.e., a temporal discounting parameter), and 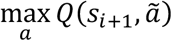 is the maximum expected value over all actions *ã* afforded in the next state *s*_*i*+1_. We defined the trials of the PRP task as consisting of the set of *i* ={1, 2, 3 , 4} event time points (1: options presented; 2: action taken; 3: outcome presented; 4: (terminal) transition screen). We modeled participant choices (*choice*_*t*_) on each trial *t* of the PRP task with a softmax choice policy (i.e., categorical logit choice model) that assigns probability to choosing each of the two options presented on a trial according to the learned Q-values of the two options present. For example, for a trial that presents option 2 and option 5, the corresponding Q-values *Q*(*s*_1_, opt_2) and *Q*(*s*_1_, opt_5) are used to compute the probability of selecting each option (e.g., option 2):

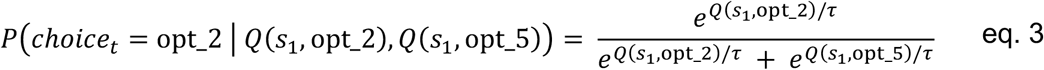

where 0 <*τ*< 20 is a choice temperature parameter that determines the softmax function slope and parameterizes an exploration versus exploitation trade-off where higher temperature values lead to a more uncertain distribution of choices and low temperature values allow choices to be attracted to higher expected values.

### Valence-Partitioned Reinforcement Learning (VPRL) model

For Valence-Partitioned RL (VPRL; Kishida and Sands, 2021), we extend the TDRL framework, but separate ‘outcomes’ and how they are processed based on the valence of the input. VPRL treats ‘Positive’ (*P*) and ‘Negative’ (*N*) input as though separate, parallel, *P*- and *N*-systems maintain a partition between appetitive and aversive input throughout processing. *P*- and *N*-system Q-values are estimated (*Q*^*p*^, *Q*^*N*^, respectively) independently, though each in a TDRL manner (see **eq. 4-7**). We model their integration in the simplest manner (**eq. 8**) when value-based decisions must be made (Note: alternative approaches for integrating these value estimates may be investigated in future work).

*P-* and *N-*systems update via TD-prediction errors on every episode, but by valence specific rules (*P-system:* **eq. 4** and *N-system:* **eq. 5)**. The *P-*system only tracks rewarding (i.e., appetitive) outcomes (*outcome*_*i*_ > 0, **eq. 4**) and the *N-*system only tracks punishing (i.e., aversive) outcomes (*outcome*_*i*_<0 , **eq. 5**); both systems encode the opposite-valence outcomes and null outcomes similarly – as though no outcome occurred.

For the *P*-system, the reward-oriented TD prediction error therefore is

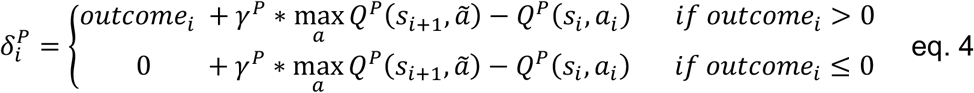

where 0 <*γ*^*p*^ < 1 is the *P-*system specific temporal discounting parameter (directly analogous to the standard TDRL temporal discounting parameter).

The *N-*system similarly encodes a punishment-oriented TD prediction error term:

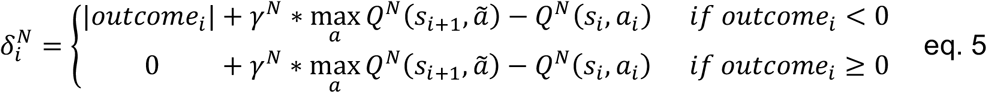

where 0 <*γ*^*N*^ < 1 is the *N-*system temporal discounting parameter and |*outcome*_*i*_| indicates the absolute value of the outcome. The absolute value of the outcome is taken to be interpreted as though the system only updates on aversive stimuli and does so based solely on the varying magnitudes.

The *P-* and *N-*systems prediction errors update expectations of future rewards or punishments of an action, respectively, according to the standard TD-learning update rule but, again, for each system independently:

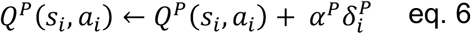

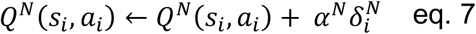

where 0 <*α*^*p*^ < 1 and 0 <*α*^*N*^ < 1 are learning rates for the *P-* and *N-*systems, *Q*^*p*^(*s*_*i*_, *a*_*i*_) is the expected state-action value learned by the *P-*system, and *Q*^*N*^(*s*_*i*_, *a*_*i*_) is the expected state-action value learned by the *N-*system.

We compute a composite state-action value term for each action by contrasting the *P-* and *N-* system Q-values,

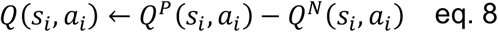

which is entered into the categorical logistic choice model (e.g., softmax policy, eq. 3) as for the TDRL model above.

### TDRL and VPRL hierarchical model parameterization

We specified a hierarchical structure to the TDRL and VPRL computational models to fit participant choice behavior on the PRP task. Individual-level parameter values (e.g., learning rates) are drawn from group-level distributions over each model parameter. This hierarchical modeling approach accounts for dependencies between model parameters and biases individual-level parameter estimates towards the group-level mean, thereby increasing reliability and certainty in parameter estimates, improving model identifiability, and avoiding overfitting (Ahn et al., 2017). These hierarchical models therefore cast individual participant parameter values as deviations from a group mean.

Formally, the joint posterior distribution *p*(*ϕ, θ*|*y, M*) over group-level parameters *ϕ* and individual-level parameters *θ* for a given model *M* conditioned on the data from the cohort of participants *y* takes the form

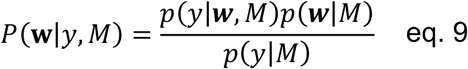

We simplify our notation to *P*(**w**|*y, M*), where **w** = {*ϕ, θ*}); here, *P*(*y*|**w**, *M*) is the likelihood of choice data *y* conditioned on the model parameters and hyperparameters, *P*(*y*|*M*) is the marginal likelihood (model evidence) of the data given a model, and *P*(**w**|*M*) is the joint prior distribution over model parameters as defined by the model, which can be decomposed into the product of the prior on individual-level model parameters conditioned on the model hyper-parameters *P*(*θ*|*ϕ, M*) times the prior over hyper-parameters *P*(*ϕ*|*M*). We define the prior distributions for individual-level model parameters (e.g., *θ*_*TDRL*_ = {*α, τ, γ*} for *M* = TDRL) and the hyper-priors of the means −∞< *µ*_(.)_< +∞ and standard deviations 0 <*σ*_(.)_< +∞ of the population-level parameter distributions (e.g., ϕ_*TDRL*_ = {*µ*_*α*_, *µ*_τ_, *µ*_*γ*_, *σ*_*α*_, *σ*_τ_, *σ*_*γ*_}) to be standard normal distributions. We estimated all parameters in unconstrained space (e.g., −∞< *µ*_*γ*_< +∞) and use the inverse Probit transform to map bounded parameters from unconstrained space to the unit interval [o, 1] before scaling estimates by the parameter’s upper bound:

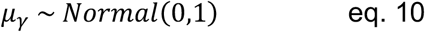

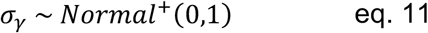

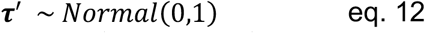

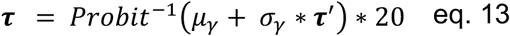

where bold terms indicate a vector of parameter values over participants. This non-centered parameterization (Papaspiliopoulos et al., 2007) and inverse Probit transformation creates a uniform prior distribution over individual-level model parameters between specified lower and upper bounds. Note that for learning rate and temporal discount parameters, the scaling factor (upper bound) was set to 1, whereas it was set to 20 for the choice temperature parameter. We used the Hamiltonian Monte Carlo (HMC) sampling algorithm in the probabilistic programming language Stan via the R package *rstan* (v. 2.21.2; Carpenter et al., 2017) to estimate the joint posterior distribution over group- and individual-level model parameters for the TDRL and VPRL models for both cohorts individually. For both models and each cohort, we executed 12,000 total iterations (2,000 warm-up) on each of 3 chains for a total of 30,000 posterior samples per model parameter. We inspected chains for convergence by verifying sufficient chain mixing according to the Gelman-Rubin statistic 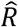, which was approximately 1 for all parameters.

### TDRL and VPRL model comparison

We compared the TDRL and VPRL models’ fits to participant choice behavior on the PRP task according to their model evidence (i.e., model marginal likelihood), which represents the probability or ‘plausibility’ of observing the actual PRP task data under each model (Mckay 2013). In Bayesian model comparison, the model with the greatest posterior model probability *P*(*M*|*y*) is deemed the best explanation for the data *y* and is computed by:

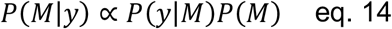

where *P*(*y*|*M*) is the model marginal likelihood (“model evidence”) and *P*(*M*) is the model’s prior probability. The model evidence is defined as:

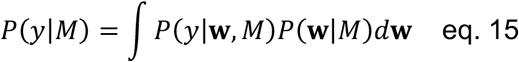

where *P*(**w**|*M*) is the prior probability of a model *M*’s parameters **w** before observing any data and *P*(*y*|**w**, *M*) is the likelihood of data *y* given a model and its parameters. We adopt the approach of approximating this integral using importance sampling (i.e., bridge sampling). Given that we only wanted to compare the TDRL and VPRL models, the relative posterior model probability can be defined as:

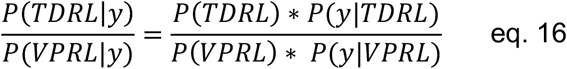

where the ratio of posterior model probabilities *P*(*TDRL*|*y*)*/P*(*VPRL*|*y*) is referred to as the “posterior odds” of TDRL relative to VPRL; *P*(*TDRL*) and *P*(*VPRL*) are the prior probabilities of the TDRL and VPRL models, respectively; and the ratio of marginal likelihoods *P*(*y*|*TDRL*)*/P*(*y*|*VPRL*) is termed the “Bayes factor”, which is a standard measure for Bayesian model comparison. Granting equal prior probabilities over the set of candidate models, each model’s evidence *P*(*y*|*M*) can be used to rank each model in the set for comparison. The marginal likelihoods are computed as log-scaled. We estimated the log model evidence for the TDRL and VPRL models for each cohort using an adaptive importance sampling routing called bridge sampling as implemented in the R package *bridgesampling* (v. 1.1-2; Gronau et al., 2017). Bridge sampling is an efficient and accurate approach to calculating normalizing constants like the marginal likelihood of models even with hierarchical structure and for reinforcement learning models in particular (Gronau et al., 2017). To further ensure stability in the bridge sampler’s estimates of model evidence, we performed 10 repetitions of the sampler and report the median and interquartile range of the estimates of model evidence. The model with the maximum (i.e., less negative) model evidence is preferred, and therefore a positive value for the difference between the log model evidences for TDRL and VPRL (as reported in **Table 1**) favors TDRL, while a negative value favors VPRL.

In addition to the standard Bayesian model comparison using model marginal likelihoods, we estimated each model’s Bayesian leave-one-out (LOO) cross-validation predictive accuracy, defined as a model’s expected log predictive density (ELPD-LOO):

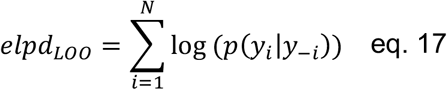

where the posterior predictive distribution *p*(*y*_*i*_|*y*_−*i*_) for held-out data *y*_*i*_ given a set of training data *y*_−*i*_, is

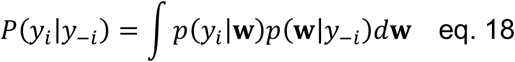

The ELPD is an estimate of (i.e., approximation to) the cross-validated accuracy of the TDRL or VPRL models in predicting new (i.e., held-out) participant data, given the posterior distribution over model parameters fit to a training set of participant data (Vehtari et al., 2017). Again, we approximate this integral via importance sampling of the joint posterior parameter distribution given the training data *p*(**w**|*y*_−*i*_). Furthermore, the upper tail of the distribution of importance weights are smoothed by a Pareto distribution (Pareto-smoothed importance sampling, PSIS) to improve the ELPD-LOO estimation. We calculated the model ELPD in this way using the R package *loo* (v. 2.3.1; Vehtari et al., 2017).

### Subjective rating computational modeling and cross-validated Bayesian regression analysis

We defined a Bayesian linear regression model of Positive and Negative valence system prediction errors and estimated Q-values on participants’ self-reported subjective feelings about their most recent outcomes measured throughout the PRP task (query probability = .33 on each trial). We express the subjective rating on a trial as normally distributed with a mean *E*(*y*_*i*_|*β, X*) that is a linear function of Positive and Negative system prediction errors and learned Q-values (predictor variable matrix *X*):

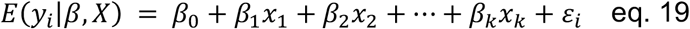

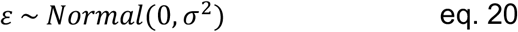

where *E*(*y*_*i*_) is the subjective rating (*i* = 1 … 50 indexes the numbers of ratings) from a participant, *β*_*k*_ are the *k* = 7 linear model weights, *X*_*k*_ are rows of the predictor variable matrix *X* (corresponding to each trial on which a subjective rating was sampled), and ε_*I*_are the normally-distributed errors with variance *σ*^2^. We define *θ*= {*β*_0_, *β*_1_, … , *β*_*k*_, *σ*} as the vector of all model parameters. The Bayesian rendering of the subjective rating linear regression model is therefore

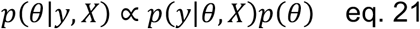

where *p*(*y*|*θ, X*) is the (normally-distributed) data likelihood function and *p*(*θ*) = *p*(*β*)*p*(*σ*^2^) are the (weakly-informative) prior distributions over model parameters:

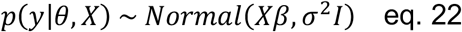

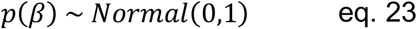

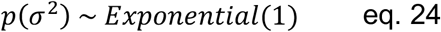

where *I* is the *n* × *n* (*n* = number of participants in training sample) identity matrix. The joint posterior distribution over model parameters *θ*conditioned on the subjective ratings and predictor variable matrix factorizes into:

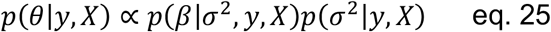

where the conditional posterior distribution *p*(*β*|*σ*^2^, *y, X*) of linear model parameters *β* conditional on *σ*^2^ is the normal distribution

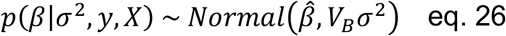

and, from the least-squares solution,

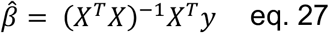

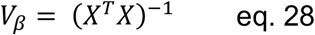

The marginal posterior distribution *p*(*σ*^2^|*y, X*) is defined as

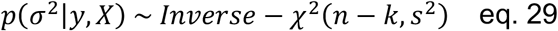

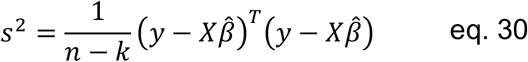

where *n*− *k* is the number of degrees of freedom (data points). We implemented this Bayesian regression model using the R package *rstanarm* (v. 2.21.1; Gabry and Goodrich, 2017), which uses HMC via Stan to efficiently sample the entire joint posterior distribution over model parameters *p*(*θ*|*y, X*). We adopted a leave-one-participant-out cross-validation approach by fitting the subjective rating regression model to all participants except for one person and drawing samples of (*β, σ*) from this fitted model’s joint posterior distribution to form a posterior predictive distribution 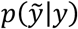 for the held-out participant’s ratings 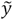 as:

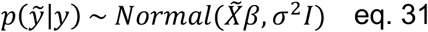

where 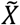 is the held-out participant’s predictor matrix. For sampling both the linear model joint posterior distribution *p*(*θ*|*y, X*) and the posterior predictive distribution 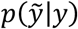, we drew 3,500 total samples (1,000 warm-up) on each of 4 chains for a total of 10,000 samples for each parameter and verified sufficient mixing according to 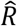 values, which were approximately 1 for all parameters.

### Functional MRI data acquisition, pre-processing, and model-based analysis

The fMRI cohort (N=47) was recruited as part of two separate studies at WFUHS, with one cohort n=20 and the other n=27. For all neuroimaging participants, we acquired fMRI and structural MRI data using a Siemens MAGNETOM 3T Skyra whole-body scanner and a 32-channel head coil. High-resolution (0.5×0.5×1.0mm^3^) T1-weighted structural MRI scans were acquired using a magnetization-prepared rapid gradient echo (MPRAGE) sequence (TR = 1480msec ; TE = 2.66msec ; flip angle = 12 degrees; FoV = 24.5cm ; 192 slices), and fMRI BOLD data were acquired by means of a multi-band (simultaneous multislice, SMS) echo-planar imaging (EPI) sequence (MB factor = 8; TR = 1000msec; TE = 30msec; flip angle = 52 degrees; FoV = 20.8cm; 72 interleaved sagittal slices; isotropic 2mm^3^ voxels). All data were pre-processed and analyzed using FSL and SPM12. Each participant’s fMRI data were aligned to a single-band reference image (SBref) and corrected for EPI (B0) distortions using a fieldmap estimated from reverse-phase encoded functional volumes (directions Right->Left and Left->Right) via FSL’s *topup* tool (Andersson et al., 2003); co-registered to the high-resolution structural volume and warped to MNI template space (2mm^3^ isotropic); spatially smoothed with a 5mm FWHM Gaussian kernel; high-pass filtered at 128 seconds (<0.008Hz); and normalized by the session grand-mean value.

For each participant, we constructed a first-level GLM to model BOLD signals during task performance. The following regressors were included in the GLM as events of interest and convolved with a canonical hemodynamic response function: (i) onset of ‘option presentation’, parametrically modulated by (a) expected value of the chosen option and (b) expected value of the unchosen option; (ii) onset of ‘outcome presentation’, parametrically modulated by the (a) ‘outcome presentation’-episode-specific positive system prediction error, (b) ‘outcome presentation’-episode-specific negative system prediction error, and (c) imputed rating of subjective feelings; and (iii) all other motor and visual stimuli. The parametric modulators at the time of outcome presentation were orthogonalized. Six head motion parameters were included as covariates of no interest. First-level GLM results for each participant were incorporated into a second-level random effects analysis at the group-level. At the group-level, all analyses were whole-brain and conducted at either a family-wise error (FWE)-corrected statistical threshold of p<0.05 or an uncorrected significance thresholds of p<1e-4 and p<1e-5.

---

**Supplemental Figure 1.**
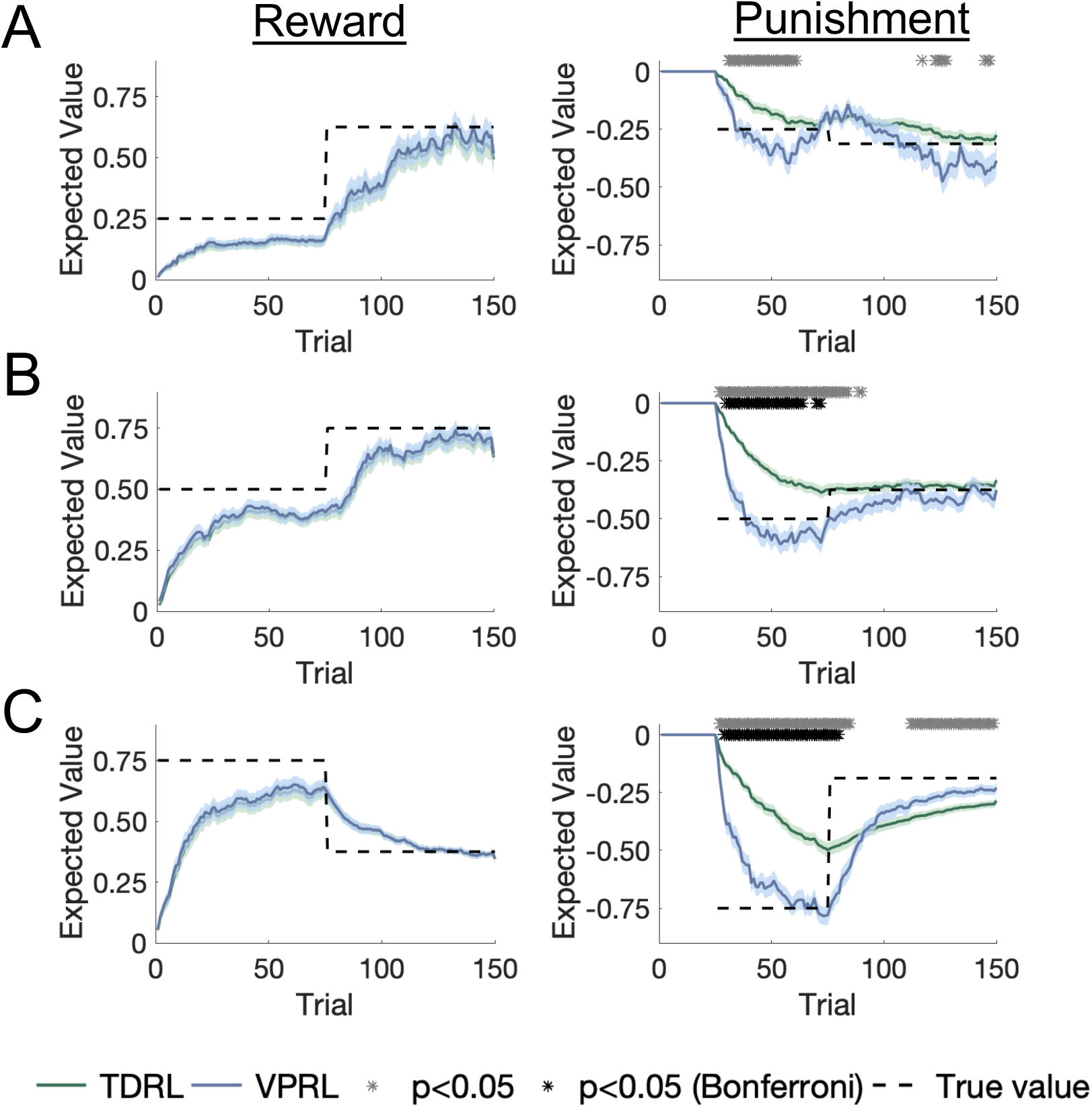
VPRL leads to more accurate learned values of punishing options on the PRP task compared to TDRL. We used each participant’s estimated parameters for the TDRL and VPRL models to compute the expected state-action value (Q-value) for each option on the PRPT task over time. The PRP options are arranged top to bottom as the (**A**) 25%, (**B**) 50%, and (**C**) 75% reward-associated (left column) or punishment-associated (right column) options. Bold green and blue traces represent mean expected value for TDRL and VPRL, respectively, and the shaded region around the means represents one standard error of the mean. TDRL and VPRL model-derived learned values for reward-associated options were very similar to each other, whereas learned values for punishment-associated options were significantly different between models, according to an independent samples t-test at each time point of the difference between the true value (dashed line) and the TDRL or VPRL model-derived learned values across participants. Grey asterisk = p<0.05, black asterisk = p<0.05 Bonferroni corrected.

**Supplemental Figure 2.**
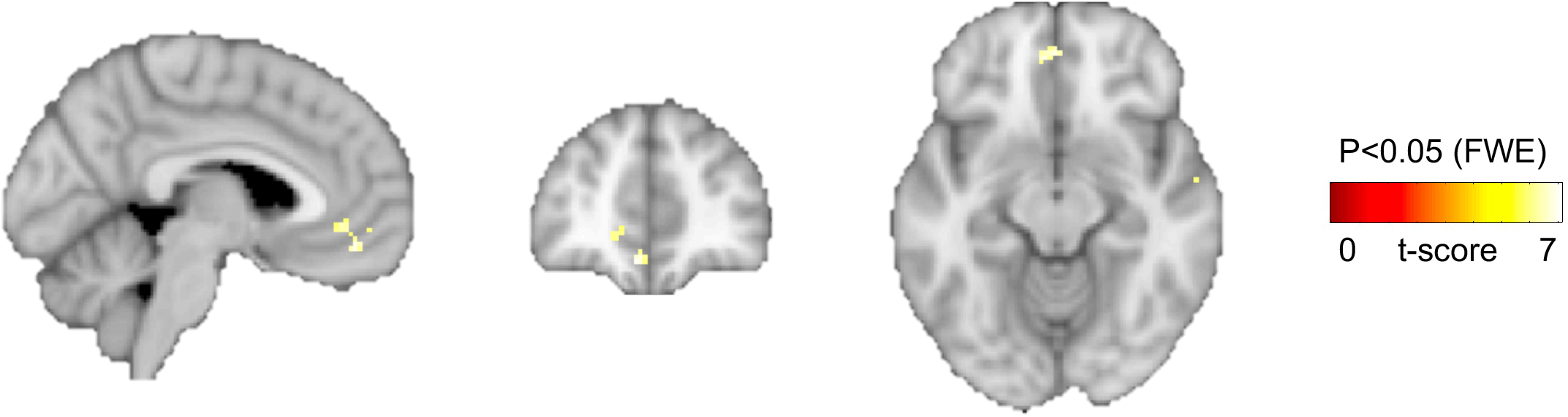
Medial prefrontal activity correlates with VPRL-derived expected state-action value signals. Whole-brain model-based analysis of VPRL learning signals indicates that medial prefrontal cortex parametrically encodes the expected values (VP-Q-value) of the chosen and unchosen options on each trial. Analyses were whole-brain FWE-corrected at p<0.05, and the slices in MNI coordinates are *X*=-4, y=46, z=-12.

